# Intake, performance, and feeding behavior of Holstein and Holstein x Gyr heifers grazing intensive managed tropical grasses during the rainy season

**DOI:** 10.1101/2021.07.13.452261

**Authors:** D. F. Quirino, M. I. Marcondes, L. N. Rennó, P. V. F. Correa, V. C. L. Morais, C. S. Cunha, T. D. A. Silva, A. L. da Silva, E. Miller-Cushon, P. P. Rotta

**Affiliations:** Department of Animal Science, Universidade Federal de Viçosa, 36570-000, Viçosa, Minas Gerais, Brasil; Department of Animal Sciences, Washington State University, 99164, Pullman, Washington, USA; Department of Veterinary Medicine, Universidade Federal de Viçosa, 36570-977 Viçosa, Minas Gerais Brazil; School of Veterinary Medicine and Animal Science, Universidade Federal de Mato Grosso do Sul, 79, Campo Grande, Mato Grosso do Sul, Brazil; Department of Animal Production, Universidade Federal Rural do Rio de Janeiro, 23897-005 20 Seropédica, Rio de Janeiro, Brazil; Department of Animal Sciences, University of Florida, 32611, Gainesville, Florida, USA

**Keywords:** bout criteria, dairy, intake, replacement, ruminating

## Abstract

Holstein × Gyr and Holstein are the primary dairy breed used in tropical systems, but when rearing under pasture, feed intake, behavior, and performance might differ between them. This study aimed to evaluate the voluntary intake, nutrient digestibility, performance, and ingestive behavior of Holstein and Holstein × Gyr (½ Holstein × ½ Gyr) heifers managed in an intermittent grazing system of Guinea grass (*Panicum maximum* Jacq. cv. Mombaça). The experiment was conducted during the summer season throughout four periods of 21 d. Two 8-heifers (four Holstein and four Holstein × Gyr) groups, averaging 258.6 ± 24.8 kg and 157.1 ± 24.99 kg body weight, were used. Each group grazed a separate set of 16 paddocks, and all heifers received a concentrate supplement daily. Heifers were weighed at the beginning and end of the experiment. Fecal, forage and concentrate samples were evaluated for their dry matter (DM), crude protein (CP), crude fat, ash, neutral detergent fiber (NDF), and indigestible NDF. Feeding behavior was evaluated through 24 h of live observation for 48 h of each experimental period. Grazing, ruminating, resting, and intake of concentrate times were recorded, and rumination criteria, bout criteria, mealtime, meal frequency, and meal duration were estimated. There was no difference in dry matter intake (DMI). The Holstein × Gyr heifers had greater NDF intake and average daily gain (ADG), and feed efficiency tended to show greater CP and NDF digestibilities. The forage DMI of Holstein × Gyr was 11.70% greater than the Holstein heifers. Holstein grazed less than Holstein × Gyr heifers in the afternoon. Ruminating time was 18.43% lower for Holstein than Holstein × Gyr heifers, and rumination criteria were greater for Holstein heifers. Holstein heifers presented more prolonged rumination bouts and resting time than Holstein × Gyr heifers. Holstein × Gyr can ingest and ruminate greater amounts of fibrous material. Holstein heifers select lower fiber material, and they need to spend more time ruminating small portions of feed. Overall, we do not recommend using young Holstein heifers in tropical pasture conditions because their ADG is low because of its lower adaptability to fibrous feed and heat stress. However, this management condition is appropriate for Holstein × Gyr heifers and results in an adequate performance.

**Implications:** This study was the first to evaluate the performance and behavior of young Holstein × Gyr and Holsteins heifers in tropical grazing systems under the same nutritional and environmental conditions. Crossbreed and purebred heifers interacted differently with the pasture; however, without noticeable variation in grazing time. As expected, Holstein heifers’ performance in the tropical pasture was impaired by a reduction in intake and grazing time. The greater performance observed for Holstein × Gyr heifers was assigned to greater forage intake, rumination time, and efficient forage nutrient use, showing animal’s adaptability to management conditions.

## Introduction

Dairy heifers are important for maintaining farms’ productivity due to their role as replacements for lactating cows (Machado et al., 2019). The performance of young dairy cattle is influenced by the quantity and quality of the feed provided and by the environmental conditions and genetics (Garcia et al., 2011). In tropical regions, pastures are used as the primary feed source due to low costs compared with intensive systems and diversity of species and forage production potential. However, pasture-based systems are often linked to low productivity, and improving these systems is of utmost importance (Macdonald et al., 2017; Machado et al., 2019).

In tropical conditions, high solar irradiation, air humidity, and temperature are commonly associated with inadequate feed management and may impair animal welfare and production. Additionally, the interaction between environmental conditions and grazing animals may cause behavior changes (Schütz et al., 2010). For example, animals in grazing systems commonly increase the search for shade and water and reduce feed appetite during the hottest hours of the day (Pereira et al., 2017) consequently, they frequently exhibit a low average daily gain (ADG) and an elevated age at first calving (Paciullo et al., 2011).

Holstein animals have been previously associated with low performance when grazing tropical grasses (Mathews et al., 1994; Machado et al., 2019); however, other studies have demonstrated that crosses between Holstein and Gyr animals generate animals able to perform better under grazing conditions (Paciullo et al., 2011; Signoretti et al., 2012). In addition, information on young cattle is frequently found in the literature; however, this information concern either pure breed or crossbreed animals. Thus, studies comparing Holstein × Gyr and Holstein heifers managed simultaneously in tropical grazing systems are still scarce.

Studies on the grazing behavior of Holstein heifers in tropical environments were not conducted before. To the best of our knowledge, this is the first study examining young Holstein and Holstein × Gyr heifers to evaluate animal performance and ingestive behavior. Reliable records are necessary to correlate heifer’s ingestive behavior and nutrient intake and digestibility, contributing to understanding the causes of change in intake profile and its impact on performance. This insight will allow the optimization of management techniques to express the maximum productive potential of animals (Santos et al., 2020).

We hypothesized that Holstein × Gyr heifers would present greater nutrient intake and greater ADG due to the better grazing ability than Holstein heifers. Therefore, this study aimed to evaluate nutrient intake, digestibility, performance, and ingestive behavior of Holstein and Holstein × Gyr heifers managed in an intermittent grazing system of Guinea grass (*Panicum maximum* Jacq. cv. Mombaça).

## Materials and methods

### Experimental design and animal management

The study was conducted at the Universidade Federal de Viçosa (Viçosa, MG, Brazil). Viçosa is located at Southwest Minas Gerais, Brazil, and the experiment was conducted during the summer season. The animals underwent a period of adaptation to the diet and management during the 45 d before the experiment. The experimental period lasted 84 d with four periods of 21 d each for data collection and sampling.

Sixteen heifers were divided into two groups (blocks) according to their initial body weight (BW) and assigned in a randomized block design with a repeated measures scheme. The average BW of the two groups was 258.6 ± 24.8 kg and 157.1 ± 24.99 kg. Each group was composed of eight heifers with four Holstein and four Holstein × Gyr (½ Holstein × ½ Gyr; Girolando), and each eight-animal block grazed a separate set of 16 paddocks. The blocks were considered the combination of the set of paddocks and the animals’ weight, and treatments were randomized within each block.

The animals were managed in a rotational grazing system comprising 32 Guinea grass (*Panicum maximum* Jacq. cv. Mombaça) paddocks with an 800 m^2^ area for a one d grazing period. The paddocks were fertilized with 200 kg of N/ha/yr and 150 kg of K_2_O/ha/yr. The animals had access to a feeding area with a feed bunk for supplementation, drinking fountains, and at least 4 m^2^ of shade per animal. Separate paddocks and feeding areas were used for each heifer group to avoid competition during concentrate supplementation. Animals were conducted daily to a new paddock after the supplement consumption (i.e., bulk empty) and stayed there until the following day.

The concentrate was provided to each heifers group at 12:00 h. The supplement was composed of soybean meal (22%), cornmeal (75%), urea (2.7%), and ammonium sulfate (0.3%) and was offered at 0.5% BW on a DM basis, updated based on animal weight every 21 d. In addition, mineral salt was provided ad libitum in the feeding area. Concentrate samples were collected directly from the feed factory before mixing the ingredients.

### Nutrient intake, digestibility, and animal performance

We used chromic oxide to estimate fecal excretion. The marker was infused (packaged in paper) at 10:00 h directly into the animal’s esophagus with the aid of a probe at a dose of 15 g per animal/d (Ribeiro et al., 2018). Individual concentrate intake was estimated using titanium dioxide as a marker. The TiO_2_ was mixed into the concentrate immediately prior to delivery at a dose of 10 g of titanium dioxide per animal/d. Lastly, we used indigestible NDF (iNDF) as an internal marker to estimate forage intake (Detmann et al., 2001). Both chromium oxide infusion and titanium dioxide supply were conducted for eight days in each period (from d -11 to d -18). Spot fecal samples were taken during the last three days of infusion (from d -16 to d – 18) in each period at 06:00, 12:00, and 18:00 h, resulting in nine samples per animal. Those samples were pooled to compose one sample per animal per period.

The grazing stratum composition and availability were determined using two isolation cages (1.0 × 1.5 m) placed in the paddocks before the animals’ entrance (from d- 16 to d- 18), in a location representing the height and density of pasture. On the day following the animals’ removal from the paddock, forage samples were taken from the cage by cutting the stratum at the same height as the pasture intake (Brandao et al., 2018).

Heifers were weighed on three consecutive days in the beginning and the end of the experiment to determine animal performance. Animals were fasted for 12 h before each weighing, with free access to water. As the concentrate supply was based on the animal’s ADG, on the first day of each period, intermediate weighings were conducted without fasting to concentrate supply and to determine the dry matter, forage, and fiber intakes as % of BW.

### Feeding behavior

Feeding behavior was evaluated through 24 h of live observation for two consecutive days (48 h) from d- 8 to d- 9 of each experimental period. We used instantaneous recording at 10 min intervals (Martin and Bateson, 2007), and the following categories were recorded: grazing, ruminating, resting, and concentrate intake. Grazing behavior was defined as the animal ingesting or selecting forage. The rumination activity occurred when the cud was being rechewing without feed ingestion. Resting behavior was defined as the nonoccurrence of ruminating nor feeding activities and happened when the animal was awake (standing or lying down) or sleeping. Lastly, the concentrate intake activity was defined as the animal eating from the bunker.

Meals were represented as eating events with short intervals within a meal, and the bout criteria were defined as the longest nonfeeding interval within a meal (Bailey et al., 2012). The time from the beginning of the feeding event until a gap between events defined the meal duration (Kargar et al., 2018).

We used the methodology described by DeVries et al.(2003) to estimate bout criteria, meal time, meal frequency, and meal duration. Briefly, the intervals between events (grazing or ruminating) were calculated and log10 transformed in Excel. The bout criteria were determined per animal per period as the point at which the distribution curve of intrameal intervals is intersected by the distribution curve of intermeal intervals (Figure 1). To estimate the bout criteria, we used the statistical software R (ver. 2.13; R Foundation for Statistical Computing) and the MIXDIST R package (http://www.math.mcmaster.ca/peter/mix/mix.html) as described by Bailey et al. (2012). Data from the bout criteria were used to calculate meal frequency (meals/d) by counting the interval numbers that exceeded the criterion plus one. The meal duration (min/meal) was calculated from the first feeding bout until the end of the last feeding bout. Finally, the total mealtime was calculated from the sum of the two meal durations described above.

**Figure 1.**
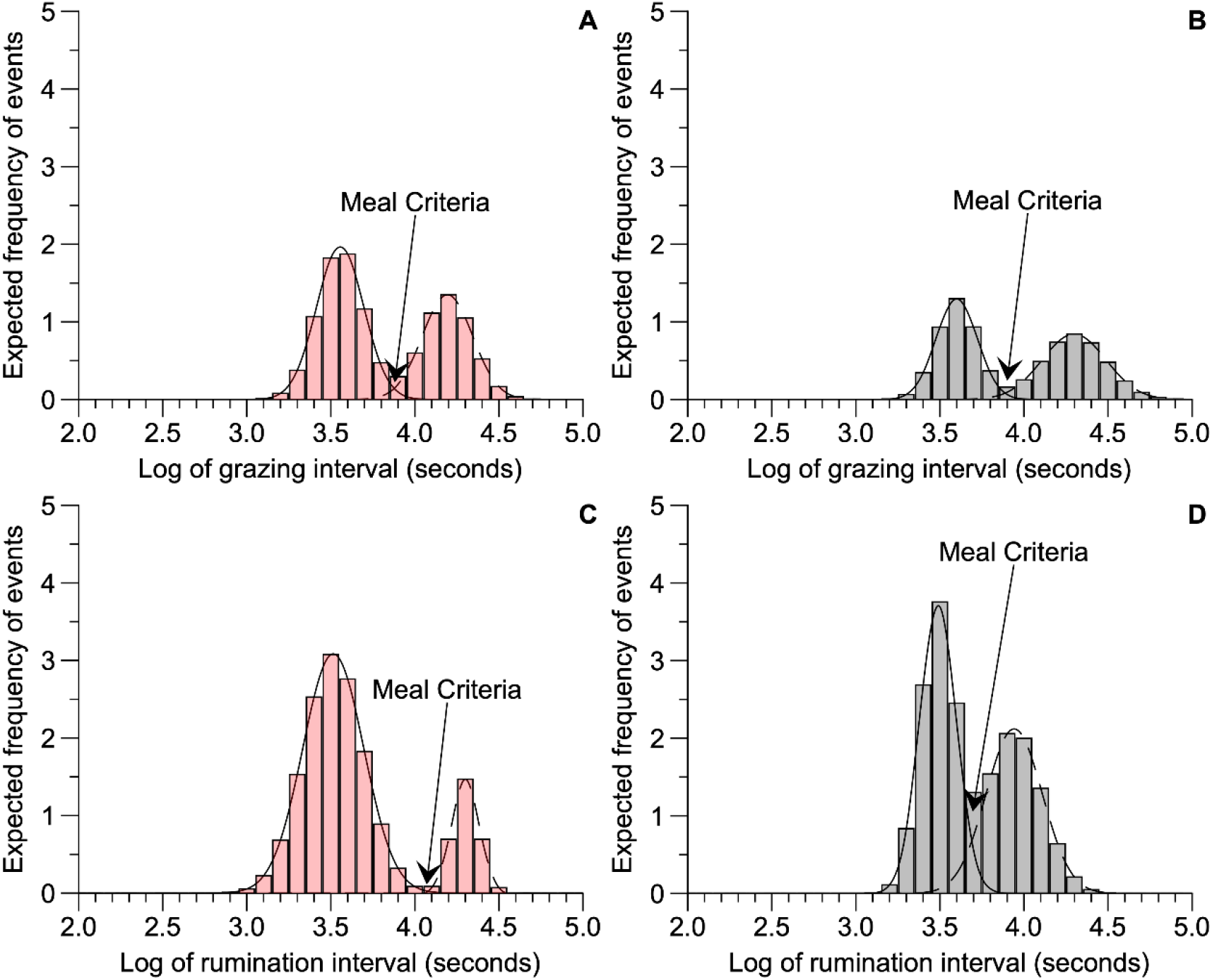
Bout criteria. The intersection of distribution intrameal curve intervals and distribution curve of intermeal intervals. **A.** Bout criteria of Holstein (red bars) heifers in grazing behavior; **B**. Bout criteria of Holstein × Gyr (grey bars) heifers in grazing behavior; **C**. Rumination criteria of Holstein (red bars) heifers in ruminating behavior; **D.** Rumination criteria of Holstein × Gyr (grey bars) heifers in ruminating behavior

### Laboratory analyses and environmental measurements

Forage and fecal samples were dried partially in a forced-air drying oven at 55°C for 72 h. Concentrate, forage, and fecal samples were ground in a Willey mill (model TE-680, brand TECNAL, Piracicaba, São Paulo, Brazil) using the 2 mm screen for iNDF analysis (INCT-CA method F-008/1) and the 1 mm screen for DM (INCT-CA G-003/1 method), CP (INCT-CA N-001/1 method), ether extract (EE) (INCT-CA G-004/1 method), ash (INCT-CA M-001/1 method), and NDF (INCT-CA F-001/1 method) analyses. Additionally, fecal samples were analyzed for titanium dioxide (INCT-CA M-007/1) as described by Detmann et al. (2012). The total digestible nutrient (TDN) content was estimated following Weiss (1999). The digestible energy (DE) in the diet was obtained by multiplying the digestible fraction of each component by its respective caloric value (ARC, 1980), and the metabolizable energy (ME) was estimated using the equation described by Galyean et al. (2016).

Data on weather conditions (average, maximum, and minimum temperatures, humidity, and rainfall) were obtained from the Viçosa Weather Station (Table 1). The temperature-humidity index (THI) was calculated using the average values of temperature and humidity, as described by Thom (1959):

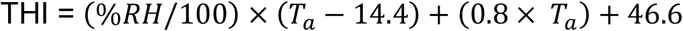

in which: THI = temperature-humidity index, %RH = relative humidity, and Ta = average air temperature in °C.

**Table 1.** Environmental conditions throughout periods

| Item | Period |  |  |  |
| --- | --- | --- | --- | --- |
|  | 1<sup>1</sup> | 2<sup>2</sup> | 3<sup>3</sup> | 4<sup>4</sup> |
| Average T, °C | 23.1 | 22.6 | 21.3 | 22.9 |
| Minimum T, °C | 19.2 | 18.9 | 17.8 | 19.8 |
| Maximum T, °C | 27.5 | 29.7 | 28.2 | 31.4 |
| Rainfall, mm | 4.2 | 3.45 | 2.9 | 4.9 |
| Relative humidity, % | 79.5 | 78.5 | 75.4 | 75.4 |
| Temperature humidity index, THI | 72.0 | 71.2 | 68.8 | 71.4 |

<sup>1</sup>January 29 to February 6, 2017.
<sup>2</sup>February 2, to March 1, 2017.
<sup>3</sup>March 16 to 22, 2017.
<sup>4</sup>April 7 to 13, 2017.

### Statistical analysis

The statistical analysis was performed using the GLIMMIX procedure of SAS (SAS University Edition). Data were analyzed using a randomized block design model:

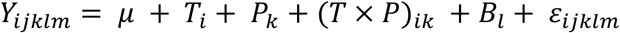

Where: μ = the general mean, *T_i_* = the fixed effect of treatment (breed) i, *P*_k_ = the fixed effect of period k, T × P_ik_ = the fixed effect of the interaction between treatment *i* and period *k, B_l_* = the random effect of block *l,* and ε_ijklm_ = random errors with a mean 0 and variance σ^2^, which is the variance between measurements within animals. Fifteen covariance structures were tested for each response variable. We used the covariance structure that provided the best fit based on a lower AIC.

To evaluate ADG and feed efficiency (FE), data that had a single observation per animal were analyzed using a randomized block design model:

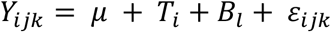

where: μ = general mean, *T_i_* = fixed effect of treatment (breed) i, *B_l_* = random effect of block *l,* and ε_ijk_ = random error with a mean 0 and variance σ^2^. Significant differences were declared when *P* < 0.05, and trends when 0.05 < *P* < 0.10.

## Results

### Forage allowance and chemical composition

The pre-grazing and post-grazing height varied from 66.44 to 74.19 cm, and from 42.40 to 45.30 cm, respectively (Table 2). In addition, we observed variation among the periods for the accumulated herbage per paddock (82.85 to 94.1 kg/DM), herbage allowance (10.3 to 11.7 kg DM/animal/d), and grazing efficiency (58.74 to 63.34%; Table 2). Forage chemical composition also changed among the periods (Table 2).

**Table 2.** Accumulated forage, pre- and post-grazing height, and chemical composition of pasture per period and supplement

| Item | Period |  |  |  | Supplement |
| --- | --- | --- | --- | --- | --- |
|  | 1 | 2 | 3 | 4 |  |
| Accumulated herbage, kg DM/ha/cycle<sup>1</sup> | 1,235.2 | 1,194.1 | 1,370.8 | 1,423.4 | - |
| Accumulated herbage, kg DM/paddock/cycle | 85.9 | 82.8 | 91.4 | 94.1 | - |
| Herbage DM allowance, kg DM/animal/cycle | 10.7 | 10.4 | 11.4 | 11.8 | - |
| Grazing efficiency, % | 61.3 | 60.1 | 58.7 | 63.3 | - |
| PreGH<sup>2</sup>, cm | 72.6 | 71.0 | 74.2 | 66.4 | - |
| PostGH<sup>3</sup>, cm | 42.4 | 43.1 | 45.3 | 44.6 | - |
| Chemical composition, %DM |  |  |  |  |  |
| Dry matter | 28.4 | 22.2 | 27.4 | 25.8 | 89.4 |
| Neutral detergent fiber | 69.3 | 70.1 | 70.3 | 66.0 | 11.0 |
| Indigestible neutral detergent fiber | 14.7 | 15.7 | 15.9 | 15.3 | 14.8 |
| Crude protein | 12.7 | 12.4 | 11.8 | 13.0 | 26.2 |
| Ether extract | 1.2 | 1.0 | 1.5 | 1.4 | 3.6 |
| Ash | 2.1 | 3.2 | 2.1 | 2.4 | 2.5 |

<sup>1</sup>Cycle of 15 d.
<sup>2</sup>PreGH = Pre-grazing height.
<sup>3</sup>PostGH = Post-grazing height.

### Nutrient intake, digestibility, and animal performance

We did not observe differences in DMI (*P* = 0.22; Table 3) between breeds, but the forage dry matter intake (FDMI) of Holstein × Gyr heifers was 11.70% greater (*P* < 0.05) than the Holstein heifers. We observed a trend (*P* = 0.07; Table 4) for greater concentrate intake (CDMI) in Holstein heifers (1.40 kg/d) compared with Holstein × Gyr heifers (1.18 kg/d). The NDF intake (NDFI) was greater (*P* < 0.05; Table 3) for Holstein × Gyr heifers compared with Holstein heifers. There was no breed effect on CP intake (CPI; *P* = 0.16), TDN intake (TNDI; *P* = 0.16) or ME intake (MEI; *P* = 0.24).

**Table 3.** Nutrient intake of grazing Holstein × Gyr and Holstein heifers

| Items | Treatment |  |  | Period |  |  |  | *P*-value |  |  |
| --- | --- | --- | --- | --- | --- | --- | --- | --- | --- | --- |
|  | Holstein×Gyr | Holstein | SEM | P1 | P2 | P3 | P4 | Breed | Per<sup>1</sup> | Breed×Per |
| Intake, kg/d |  |  |  |  |  |  |  |  |  |  |
| DM | 6.06 | 5.74 | 1.237 | 5.64<sup>b</sup> | 5.36<sup>b</sup> | 5.71<sup>b</sup> | 6.89<sup>a</sup> | 0.222 | 0.001 | 0.380 |
| FDM<sup>2</sup> | 4.87 | 4.30 | 0.941 | 4.42<sup>b</sup> | 4.17<sup>b</sup> | 4.42<sup>b</sup> | 5.35<sup>a</sup> | 0.002 | <.001 | 0.383 |
| CDM<sup>3</sup> | 1.18 | 1.40 | 0.299 | 1.20 | 1.16 | 1.28 | 1.53 | 0.07 | 0.125 | 0.617 |
| NDF | 3.49 | 3.12 | 0.693 | 3.19<sup>b</sup> | 3.04<sup>b</sup> | 3.25<sup>b</sup> | 3.74<sup>a</sup> | 0.007 | 0.002 | 0.347 |
| CP | 0.91 | 0.90 | 0.184 | 0.88<sup>b</sup> | 0.83<sup>b</sup> | 0.83<sup>b</sup> | 1.08<sup>a</sup> | 0.916 | 0.001 | 0.440 |
| TDN | 3.77 | 3.53 | 0.819 | 3.43<sup>a</sup> | 3.23<sup>b</sup> | 3.51<sup>a</sup> | 4.44<sup>a</sup> | 0.162 | <.0001 | 0.525 |
| ME, Mcal/kg DMI | 6.05 | 4.98 | 3.219 | 4.73 | 4.99 | 5.72 | 6.61 | 0.249 | 0.468 | 0.860 |
| Intake, % of BW |  |  |  |  |  |  |  |  |  |  |
| DM/BW | 2.17 | 2.31 | 0.001 | 2.35<sup>a</sup> | 2.14<sup>b</sup> | 2.10<sup>b</sup> | 2.36<sup>a</sup> | 0.052 | 0.014 | 0.111 |
| FDM/BW | 1.74 | 1.80 | 0.001 | 1.87<sup>a</sup> | 1.70<sup>b</sup> | 1.64<sup>b</sup> | 1.86<sup>a</sup> | 0.281 | 0.005 | 0.083 |
| NDF/BW | 1.25 | 1.27 | 0.001 | 1.34<sup>a</sup> | 1.22<sup>b</sup> | 1.19<sup>b</sup> | 1.27<sup>ab</sup> | 0.552 | 0.022 | 0.214 |
| Digestibility, g/g |  |  |  |  |  |  |  |  |  |  |
| DM | 0.62 | 0.62 | 0.01 | 0.60<sup>b</sup> | 0.61<sup>b</sup> | 0.62<sup>b</sup> | 0.64<sup>a</sup> | 0.286 | 0.005 | 0.642 |
| NDF | 0.65 | 0.64 | 0.01 | 0.64<sup>b</sup> | 0.6<sup>b</sup> | 0.66<sup>a</sup> | 0.65<sup>ab</sup> | 0.063 | 0.053 | 0.608 |
| CP | 0.69 | 0.68 | 0.01 | 0.66<sup>b</sup> | 0.66<sup>b</sup> | 0.68<sup>b</sup> | 0.72<sup>a</sup> | 0.051 | <.001 | 0.862 |

<sup>1</sup>Per = period.
<sup>2</sup>FDM = forage dry matter intake.
<sup>3</sup>CDM = concentrate dry matter intake.
<sup>a–b</sup>Means within a row with different letters superscripts differ *P* < 0.05.

**Table 4.** Behavior activities of grazing Holstein × Gyr and Holstein heifers

| Items | Treatments |  |  | Period |  |  |  | *P*- value |  |  |
| --- | --- | --- | --- | --- | --- | --- | --- | --- | --- | --- |
|  | Holstein × Gyr | Holstein | SEM | P1 | P2 | P3 | P4 | Breed | Per<sup>1</sup> | Breed×Per |
| Grazing characteristics |  |  |  |  |  |  |  |  |  |  |
| Grazing time, min/d | 365.2 | 354.2 | 11.32 | 348.8<sup>b</sup> | 404.7<sup>a</sup> | 335.6<sup>b</sup> | 349.7<sup>b</sup> | 0.499 | <.001 | 0.271 |
| Bout criteria, min | 118.0 | 135.5 | 0.03 | 141.9<sup>a</sup> | 132.4<sup>ab</sup> | 138.6<sup>ab</sup> | 95.9<sup>b</sup> | 0.305 | 0.026 | 0.538 |
| Mealtime, min/d | 929.8 | 877.0 | 35.40 | 927.3<sup>ab</sup> | 822.8<sup>b</sup> | 895.6<sup>ab</sup> | 967.8<sup>a</sup> | 0.299 | 0.066 | 0.138 |
| Bouts, events/d | 3.8 | 3.9 | 0.19 | 3.6<sup>b</sup> | 3.9<sup>ab</sup> | 3.8<sup>ab</sup> | 4.2<sup>a</sup> | 0.612 | 0.042 | 0.446 |
| Meal duration, min/meal | 254.6 | 230.1 | 8.97 | 258.1<sup>a</sup> | 254.7<sup>b</sup> | 394.4<sup>ab</sup> | 314.1<sup>ab</sup> | 0.062 | 0.055 | 0.667 |
| Ruminating characteristics |  |  |  |  |  |  |  |  |  |  |
| Ruminating time, min/d | 418.9 | 341.7 | 13.31 | 397.8<sup>ab</sup> | 400.0<sup>a</sup> | 364.4<sup>bc</sup> | 359.1<sup>c</sup> | <0.001 | 0.060 | 0.593 |
| Rumination criteria, min | 93.7 | 118.0 | 0.32 | 105.2 | 107.6 | 100.4 | 102.8 | 0.032 | 0.953 | 0.379 |
| Rumination total time, min/d | 719.1 | 729.2 | 26.89 | 837.2<sup>a</sup> | 696.6<sup>ab</sup> | 811.6<sup>a</sup> | 551.3<sup>b</sup> | 0.794 | 0.002 | 0.663 |
| Rumination bouts, events/d | 4.7 | 4.3 | 0.30 | 5.5<sup>a</sup> | 4.4<sup>ab</sup> | 4.8<sup>a</sup> | 3.4<sup>b</sup> | 0.371 | 0.004 | 0.631 |
| Rumination length, min/bouts | 157.1 | 179.6 | 6.94 | 164.1 | 166.2 | 178.8 | 164.3 | 0.026 | 0.674 | 0.652 |
| Other behaviors |  |  |  |  |  |  |  |  |  |  |
| Resting time, min/d | 628.8 | 712.5 | 23.0 | 662.5<sup>b</sup> | 602.5<sup>c</sup> | 714.4<sup>a</sup> | 703.1<sup>ab</sup> | <0.001 | 0.159 | 0.514 |
| Concentrate intake time, min/d | 26.7 | 31.3 | 2.77 | 30.6<sup>a</sup> | 32.5<sup>a</sup> | 24.7<sup>b</sup> | 28.12<sup>ab</sup> | 0.059 | 0.007 | 0.050 |

<sup>1</sup>Per = period.
<sup>a–b</sup>Means within a row with different letters superscripts differ $P < 0.05$ .

We observed a trend (*P* = 0.05) for greater DMI/BW (Table 3) in Holstein heifers (2.31%) compared with Holstein × Gyr heifers (2.17%). Breed did not influence (*P* = 0.28) the FDMI per BW (FDMI/BW). We observed trends for a greater NDF (*P* = 0.06) and CP (*P* = 0.05) digestibilities in Holstein × Gyr heifers than Holstein heifers (Table 3).

The final BW of Holstein × Gyr heifers was 311.50 ± 23.7 kg and the final BW of Holstein heifers was 260.39 ± 28.7 kg. The ADG (*P* = 0.01; Figure 2) and feed efficiency (*P* < 0.05; Figure 2) were greater for Holstein × Gyr heifers compared with Holstein heifers.

**Figure 2.**
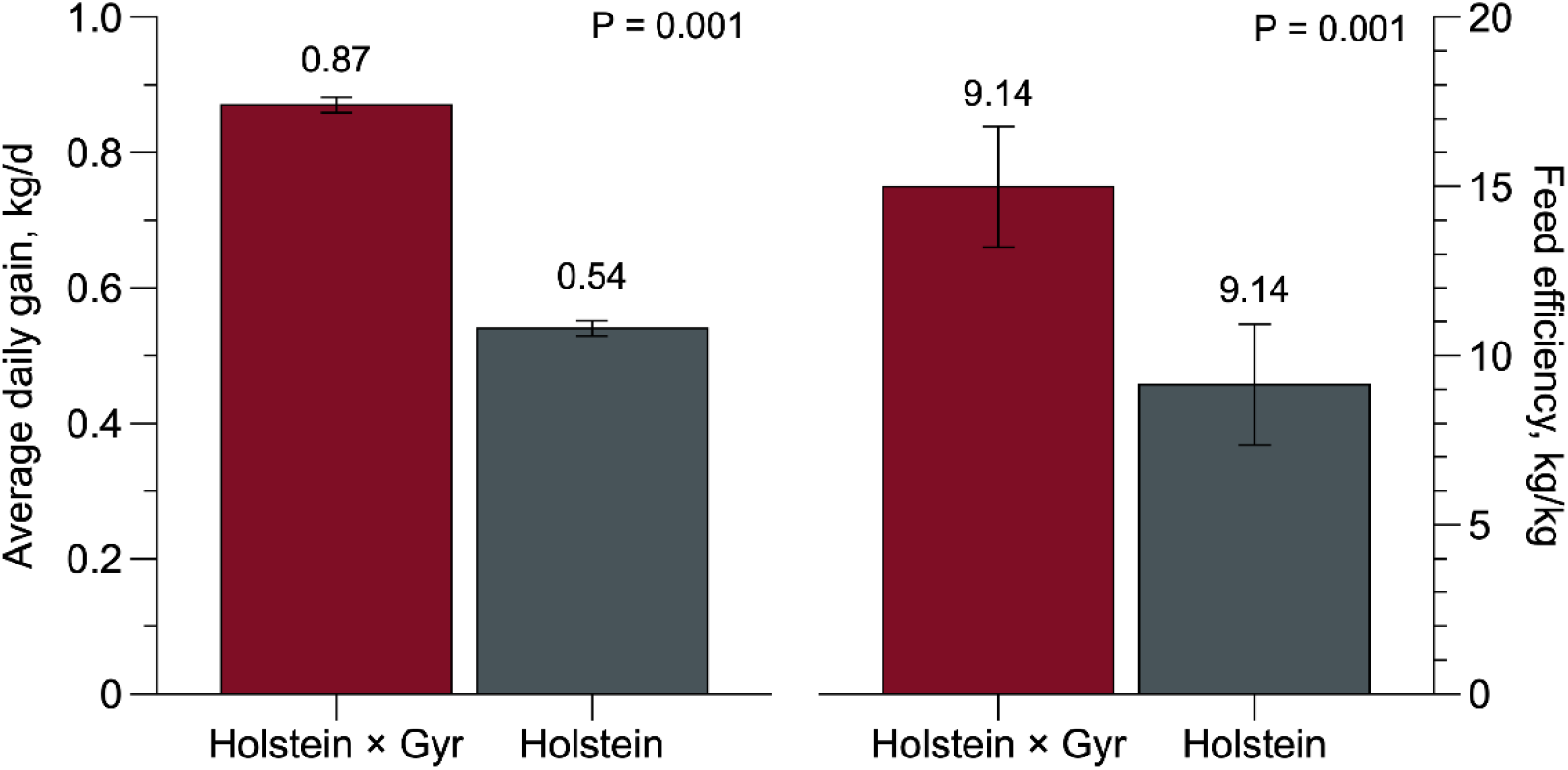
Average daily gain and feed efficiency of Holstein × Gyr and Holstein heifers grazing intesively managed Guinea grass (*Panicum maximum* Jacq. cv. Mombaça) pasture.

### Feeding behavior

The grazing time was not affected by breed (*P* = 0.49; Table 4), but grazing behavior differed. All heifers preferred to graze during the morning (05:00 to 09:00 h; Figure 3a), but a greater percentage of Holstein heifers was observed (61,3 % *v.* 47,5 %).In the afternoon (13:00 to 17:00 h; Figure 3a), the percentage of animal grazing was 10.5 % greater for Holstein × Gyr than Holstein heifers (74.6 % *v*. 64,1%; Figure 3a). In the night (19:00 to 23:00 h), a discrete difference was found, and Holstein heifers were the majority of animals grazing (30,2% *v*. 28,3%). Breed did not influence bout criteria (*P* = 0.30), mealtime (*P* = 0.29) or number of bouts events (*P* = 0.60; Table 4). A trend (*P* = 0.06) for longer meal duration in Holstein × Gyr heifers (254.63 min/d; Table 4) was observed.

**Figure 3.**
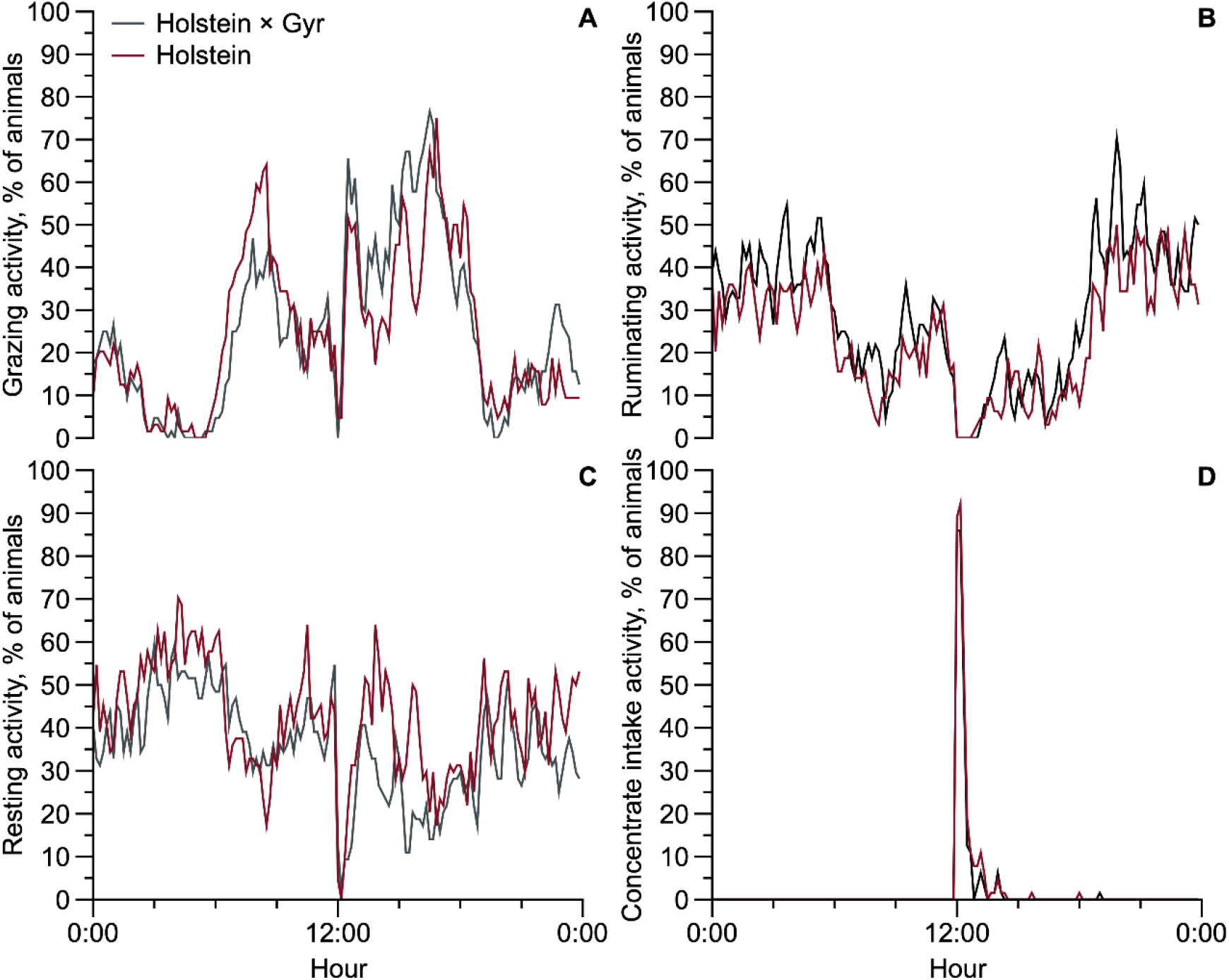
Animals’ percentage in behavior activities during 24 h grazing intesively managed Guinea grass (*Panicum maximum* Jacq. cv. Mombaça) pasture: **A**. grazing activity; **B.** ruminating activity; **C.** resting activity; **D.** concentrate intake activity. Average data from the four experimental periods

Ruminating time was affected by breed (*P* < 0.05) and was 18.43% lower for Holstein when compared with Holstein × Gyr heifers (Table 4). The rumination criteria were greater (*P* = 0.03) for Holstein than Holstein × Gyr heifers. Breed did not influence the total rumination time (*P* = 0.79) or the number of rumination bouts (*P* = 0.37). We observed longer rumination bouts (*P* = 0.02) for Holstein heifers (179 min/bouts). The resting time of Holsteins was 11.5% greater (*P* < 0.05) than that of Holstein × Gyr heifers. We observed a trend for greater concentrate intake time (*P* = 0.05) in Holstein heifers, which spent 31.25 min/d in the bunker compared with the 26.71 min/d spent by Holstein × Gyr heifers.

## Discussion

Contrary to our expectations, we did not observe breed effects on DMI, although performance was affected. This fact contradicts Santos et al. (2011), who affirmed that Holstein heifers managed in the tropical pasture have their intake compromised. As the animal metabolizable energy intake is affected by intake and digestibility (Lemaire et al., 2011), Holstein heifers probably selected the diet to achieve their nutritional requirements (Van Soest, 1994), searching for plant parts with greater CP and mineral contents and higher in digestibility (Lemaire et al., 2011). This speculation agrees with our findings where Holstein heifers had lower NDF and FDM intakes and a tendency to greater concentrate intake.

On the other hand, we observed greater FDM and NDF intakes and a trend for greater NDF digestibility for Holstein × Gyr heifers, which likely compensated for their lower concentrate intake. Furthermore, as Holstein × Gyr had a greater ADG than Hosltein, Holstein × Gyr heifers probably increased their organs feed capacity (Santana Junior et al., 2013), which had a positive effect on NDFI. As a consequence, increasing the NDFD resulted in better forage nutrient utilization. These statements reinforce the lack of difference in DMI between breeds and greater performance observed for Holstein × Gyr animals.

Holstein × Gyr and Holstein heifers interact with the grazing system differently, although no difference was observed in grazing time. As meal duration was greater for Holstein × Gyr heifers, this means they spent more time foraging or grazing (Charlton and Rutter, 2017), which supports our findings of greater FDMI and NDFI by these animals. Describing our impression as observers, when animals were conducted to a new paddock, Holstein heifers presented a reluctance to get off from the feeding area (i.e., next to the bunk feed). Additionally, these heifers had searched for palatable leaves when grazing, and this might be related to adjustment in grazing behavior to be more selective (Van Soest, 1994; Lemaire et al., 2011). Therefore, these heifers presented lower FDMI and lowered ruminating time, culminating in a lower energy intake and digestibility. In this sense, Holstein heifers increased time spent eating concentrate (or were more aggressive in the feed bunk) to achieve their daily nutrients requirements.

In contrast, Holstein × Gyr heifers usually start grazing immediately after the transfer, with less selective behavior. The greater ruminating time observed on Holstein x Gyr heifers can explain the trends for greater NDF and CP digestibilities. Thus, even though Holstein x Gyr and Holstein heifers have eaten similar quality forage, the first ones ruminated more and presented greater digestibility. Additionally, the trend of greater CP digestibility for Holstein × Gyr probably improved the N utilization by rumen microorganisms resulting in animal performance improvement (Moraes et al., 2006). Better N use and greater ability in degrading fibrous material compared to Holstein corroborate our findings, where Holstein × Gyr had feed efficiency and ADG 37% greater on average than Holstein heifers.

Ingestive behavior is intrinsically bound to feed characteristics, and different breeds might present differences in pasture intake even when grazing together (Oldenbroek and Jansen, 1978). The animal’s selective behavior leads to a bite size reduction, which searches for succulent plant parts increase the grazing time. However, there is a maximum grazing time daily available to an animal, and frequently a bite size reduction also reduces the intake (Lemaire et al., 2011). Unfortunately, we did not evaluate the bite size, but there was a reduction in forage intake for Holstein compared with Holstein × Gyr. As the animals grazed together, both breeds might be reached the maximum grazing time daily, interacting with pasture differently. As discussed previously, Holstein heifers likely spent more time selecting for leaves with lower NDF content.

Additionally, we speculate that the presence of animals from another breed (Holstein × Gyr) might have affected the grazing behavior of Holstein animals in that environment, in this case, increase grazing time, as pointed by Oldenbroek and Jansen (1978). Thus, we suspect that social interactions might play an essential role in the feeding behavior of grazing animals, and future studies are warranted. This fact might help us understand the lack of difference in grazing time between breeds.

Holstein × Gyr heifers spent a greater time on rumination activity (418.91 min/d) which is directly related to the fiber content ingested as discussed. Conversely, Holstein heifers spent less time ruminating, reinforcing the diet selection occurrence and the lower NDF ingestion. Ruminating time was 341.72 min/d for Holstein heifers and 418.91 min/d for Holstein × Gyr heifers, consistent with data ranging from 236 to 610 min/d for dairy cows observed by White et al. (2017). In addition, a greater rumination time occurred at night for both breeds (Figure 3b).

Heat stress was initially thought to be the primary influencer on the results found in the present study. There is a substantial decrease in DMI, rumination time, growth, and feed efficiency when animals are outside their thermoneutral zone (Oliveira and Ferreira, 2016). We observed a depressing rumination time for Holstein heifers and increased rumination criteria and rumination length (min/bouts) compared to Holstein × Gyr. Thus, Holstein spent more time per event ruminating less content, reflecting lower FDM and NDF intake. In hot environments, the energy spent to dissipate the heat produced from fiber fermentation increases, reducing energy utilization by animals (Beede and Collier, 1986). The heat produced by digestion needs to be dissipated generally by increasing water intake and paint rate to alleviate the heat load (Bernabucci et al., 2010). These observations combined with a lower NDFI might explain how Holstein heifers deal with the heat produced during the digestive process.

Furthermore, Holstein heifers spent more time (712.50 min/d) in resting behavior than Holstein × Gyr heifers (628.75 min/d), which is related to less time ruminating or might be associated with thermal discomfort since, in heat stress situations, animals may reduce rumination activity by increasing their search for shade (Pereira et al., 2017). The THI conditions during the experiment ranged from 68.8 to 72.0, which might cause moderate heat stress (De Rensis et al., 2015) and probably contributed to Holstein’s lower performance. However, despite adaptations to grazing activity observed in Holstein heifers (i.e., grazing occurred mainly in the early morning and late afternoon), the time spent grazing was similar between breeds, so we cannot associate all the results observed herein merely to heat stress. It is worth noting that behavioral analyses are sometimes passive to bias because of the evaluators’ presence, disturbing animals’ natural behavior. This is particularly important in the every-minute evaluations (Beauchemin, 2018). Therefore, our study aimed to avoid this interference by collecting behavioral data every 10 min (Martin and Bateson, 2007) and using a minimum distance of 10 m from the animals.

In summary, the better performance of the Holstein × Gyr animals may be related to greater forage intake, rumination time, and efficient nutrient use. While Holstein × Gyr can ingest and ruminate more significant fibrous material, Holstein heifers select lower fiber material on pasture and increase concentrate intake. Additionally, they need more time ruminating small feed portions to deal internally with dietary fiber metabolism. It is of particular importance to highlight that Holstein and Holstein x Gyr heifers grazed for a similar time, contradicting the expectations, since it was thought that Holstein would suffer severe heat stress and, thus, spend less time grazing. Therefore, there is a need for studies on other factors influencing Holstein heifers’ performance on pasture, such as ruminal parameters, rumen microbiota, tick incidence, and elucidating the fundamental role of heat stress.

## Conclusion

Overall, we do not recommend using young Holstein heifers in tropical pasture conditions due to lower ADG and its lower adaptability to fibrous feed and heat stress. However, this management condition seems appropriate for Holstein × Gyr heifers and results in adequate performance.

## Ethics approval

The experiment was performed following the standard procedures for Animal Care, and Handling stated in the Universidade Federal de Viçosa guidelines process number 24.

## Author contributions

Conceptualization: Polyana Pizzi Rotta

Methodology: Polyana Pizzi Rotta, Marcos Inácio Marcondes

Validation: Daiana Francisca Quirino, Emily Miller-Cushon, Alex Lopes Silva

Formal Analysis: Alex Lopes Silva, Marcos Inácio Marcondes

Investigation: Daiana Francisca Quirino, Valber Carlos Lima Moraes, Pietro Vítor Felix Correa,

Resources: Polyana Pizzi Rotta, Marcos Inácio Marcondes,

Data Curation: Daiana Francisca Quirino, Marcos Inácio Macondes, Alex Lopes Silva, Camila Soares Cunha

Writing – Original Draft: Polyana Pizzi Rotta, Daiana Francisca Quirino, Marcos Inácio Marcondes, Alex Lopes Silva

Writing – Review & Editing: Daiana Francisca Quirino, Camila Soares Cunha

Visualization: Polyana Pizzi Rotta, Marcos Inácio Marcondes

Supervision: Polyana Pizzi Rotta

Project Administration: Daiana Francisca Quirino, Polyana Pizzi Rotta

Funding acquisition: Polyana Pizzi Rotta

